# Short term effect of tetrahydrocurcumin on adipose angiogenesis in very high-fat diet-induced obesity mouse model

**DOI:** 10.1101/2023.05.05.539564

**Authors:** Bhornprom Yoysungnoen, Umarat Srisawat, Pritsana Piyabhan, Nattapon Sookprasert, Naphatsanan Duansak, Nakorn Mathuradavong, Natwadee Poomipark, Narongsuk Munkong, Chatchawan Changtam

## Abstract

Tetrahydrocurcumin (THC) has been shown to possess anti-angiogenic activities. This study aims to investigate the effects of THC on adipose angiogenesis and expression of angiogenic factors that occurs in 60% high-fat diet-induced obese mice. Male ICR mice were randomly divided into 3 groups: mice fed with a low-fat diet (LFD group); mice fed with very high fat diet (VHFD group), and mice fed with VHFD supplemented with THC (300 mg/kg/day orally)(VHFD+THC group) for 6 weeks. Body weight (BW), food intake, fasting blood sugar (FBS), lipid profiles and visceral fats weight (VF) were measured. The microvascular density (MVD), VEGF, MMP-2 and MMP-9 expressions were evaluated. The VHFD group had significantly increased total cholesterol, triglyceride, food intake, BW, VF, VF/BW ratio, adipocyte size and the number of crown-liked structures as compared to LFD group. THC supplementation markedly reduced these parameters and adipocyte hypertrophy and inflammation in white adipose tissues. MVD, VEGF, MMP-2 and MMP-9 were over-expressed in the VHFD group. However, THC supplementation decreased MVD and reduced expression of VEGF, MMP-2 and MMP-9. In conclusion, THC suppressed angiogenesis in adipose tissue by the downregulation of VEGF, MMP-2 and MMP-9. With its effects on lipid metabolism as well as on food consumption, THC could contribute to lower visceral fat and body weight. Overall, our study demonstrated the potential benefit of THC in mitigating obesity and associated metabolic disorders along with elucidated the suppression of adipose angiogenesis as one of its underlying mechanisms.

**Author summary:** Conceptualization, B.Y, U.S., P.P., N.D., N.S. and N.M3.; methodology, B.Y., U.S., P.P., N.D., N.S., N.M^3^., and C.C; validation, B.Y., U.S., P.P., N.D., N.S., N.M^1^., and N.M^3^.; formal analysis, B.Y., U.S, N.S., N.M^1^., N.P., and N.M^3^; investigation, B.Y., U.S, N.S., N.M^1^., N.P., and N.M^3^.; resources, B.Y. and C.C.; data curation, B.Y. and N.M^1^.; writing—original draft preparation, BY; writing—review and editing, B.Y; visualization, B.Y., U.S., P.P., and N.D.; supervision, B.Y.; project administration, B.Y., U.S., P.P., N.S., and N.P.; funding acquisition, B.Y., U.S., P.P., N.S., and N.P.

## Introduction

Obesity is a current global major health problem. Excess body weight increases the risk of several diseases, including hypertension, cardiovascular disease, cerebrovascular disease, type 2 diabetes, and cancer (1-3). Obesity is characterized by adipocyte hyperplasia and hypertrophy leading to an increase of adipose tissue mass (4). Like the growth of cancerous tumors, the growth of adipose tissue requires neo-vascularization process to supply growing adipose tissue with nutrients and oxygen. Moreover, recent data show that adipogenesis and angiogenesis are closely related during adipose tissue development (5). Therefore, the inhibition of angiogenesis in adipose tissue can potentially be a strategy to prevent adipose tissue growth and subsequent obesity.

Vascular endothelial growth factor (VEGF) and its receptor (VEGFR) are attractive targets for anti-angiogenic therapy to reduce obesity as they play an important role in adipose angiogenesis (6, 7). In expanding adipose tissue at the early stages of a High-fat diet (HFD)-induced obesity, expression of VEGF in white adipose tissue (WAT) enhances angiogenesis. A previous study demonstrated that anti-VEGF antibody inhibited not only angiogenesis, but also the formation of adipose/angiogenic cell clusters (8) indicating that VEGF is a key mediator of angiogenesis as well as adipogenesis.

Extracellular matrix (ECM) proteolysis is required for cell migration during blood vessel development and for adipose tissue expansion. Matrix metalloproteinases (MMPs), including MMP-2 and MMP-9, are key factors involved in ECM degradation, and their main actions in adipose tissues include adipogenesis, angiogenesis, and adipose tissue expansion. A previous study showed that MMP inhibitors significantly reduced gonadal and subcutaneous adipose tissue masses in HFD-fed wild-type mice (9). MMP inhibitors also reduced body weight gain in obese mice (10). Moreover, several types of angiogenesis inhibitors, such as TNP-470, and VEGFR-2 inhibitors, have been shown to inhibit adipose tissue expansion in mice (11-13), suggesting that the inhibition of these substances could reduce adipose tissue development. Overall, these findings have provided more information about the possible therapeutic interventions of obesity and obesity-associated disorders by targeting the vascular compartment.

The use of anti-angiogenic herbal medicines for the regulation of adipose tissue growth has gained interest owing to their safety and efficacy in treating obesity. Among the phytochemicals studies, researchers pay more attention to polyphenols, which are derived from natural plants. One such plant is the root of *Curcuma longa* L. or turmeric, which is widely cultivated mainly in tropical regions of Asia and has been consumed daily without reported toxicity (14). Curcumin, the major polyphenol in turmeric spice, showed anti-obesity in both *in vitro* and *in vivo* models (15-17). Interestingly, supplementation with curcumin exerts anti-obesity effect through suppression of angiogenesis in mice fed a high-fat diet (15). However, curcumin is poorly absorbed from the gastrointestinal tract and undergoes biotransformation by intestinal bacteria (18-20). Therefore, curcumin metabolites have been developed. Tetrahydrocurcumin (THC), the main metabolite of curcumin, lacks α, β-unsaturated carbonyl moiety and is white in color and stable in phosphate buffer as well as in saline at various pH values, making it different from curcumin (21). Moreover, THC is easily absorbed through the gastrointestinal tract, suggesting that THC might play a crucial role in curcumin-induced biological effects. THC has been shown to have anti-oxidation (22, 23), anti-inflammation (24, 25), and anti-cancer activities (25-27). Our previous studies also found that THC possesses anti-angiogenic activities in tumor (28, 29) which was more potent than that of curcumin (29). The effect of THC on angiogenesis in adipose tissue in very high-fat diet-induced obesity mouse model, however, has not been reported. Because THC is known for its antiangiogenic activity and its ability to suppress tumor growth, we hypothesized that THC could prevent adipose tissue growth via the inhibition of angiogenesis. The current study thus investigated the effects of THC supplementation on adipose angiogenesis and on the expression of angiogenic factors in very-high-fat diet-induced obese mice.

## Materials and methods

### Animals and Experimental Model

All experimental protocols were reviewed and approved by the Animal Ethics Committee of Thammasat University (approval code 020/2021). Male ICR mice (20–25 g) were purchased for the Siam Nomura International Co. Ltd. and were transported to the Animal Laboratory Center of Thammasat University. The animals were acclimatized under standard laboratory conditions, and allowed access to food and water *ad libitum* for one week. After 1 week of acclimation, the mice were divided into 3 groups: 1) low-fat control diet (LFD) group (n = 8); mice were fed of LFD (7% kcal from fat, CP082G, National laboratory Animal Center, Thailand); 2) Very high-fat diet (VHFD) group (n = 8); mice were fed of VHFD (60% kcal from fat, MP Biomedicals, USA) and treated with distilled water for 6 weeks (VHFD+vehicle group); 3) VHFD+THC Treated group (n = 8); VHFD-fed mice were treated with 300 mg/kg of THC. All treatments were given orally every day for 6 weeks. The body weight and energy consumption were recorded weekly. At the end of the treatment, mice were overnight fast and were anesthetized prior to blood collection via cardiac puncture. The serum was prepared for laboratory analyses including fasting blood glucose and lipid profiles. Their visceral white adipose tissues (vWATs) from retroperitoneal, and mesenteric regions were collected and weighed. The total vWATs weight of each mouse was recorded. The relative adipose tissue weight is expressed as the total vWATs per final body weight of each mouse. vWATs were fixed in 10% formalin for further analyses.

### Histological analysis

Paraffin embedded vWATs were stained with hematoxylin and eosin (H&E). To determine the average adipocyte size, 100 adipocytes/mouse were measured in 10 random ×10 microscopic fields from 8 mice per group, using the Image J 1.38 software (National Institutes of Health, USA). The number of CLSs was measured using a protocol as previously described (30).

### Immunohistochemistry for CD31 expression and Microvascular density (MVD) determination

To quantify angiogenesis, microvascular density (MVD) was assessed by immunostaining with the anti-CD31 antibody. The vWAT samples were incubated with primary monoclonal antibody CD31 (Ready to use, DAKO cytomation, USA) followed the protocol described previously (28). The sections were observed first under the low power (×40), then the densest area of microvessel sections was selected and captured 3-5 independent fields of view per mouse. The percentage of the CD31 immunoreactivated area to the total area was analyzed by ImageJ 1.38 software (National Institutes of Health, USA).

### Immunohistochemistry for angiogenic biomarkers

Paraffin sections of vWATs were incubated with primary monoclonal antibody VEGF (Ready to use, DAKO cytomation, USA), VEGFR-2 (1:200; ab115805, Abcam, USA), MMP-2 (1:500; ab86607, Abcam), MMP-9 (1:500; ab288402, Abcam, USA) at 4 °C for one hour. After rinsed with PBS, the samples were developed by the Envision system/HRP (DAKO cytomation, USA) for 30 min and substrate-chromogen for 10 min at room temperature. The percentage of the VEGF, VEGFR-2, MMP-2, and MMP-9 immunoreactivated area to the total area was calculated using ImageJ 1.38 software (National Institutes of Health, USA).

### Serum analysis

The concentrations of serum glucose, triglyceride (TG), total-cholesterol, low-density lipoprotein cholesterol (LDL-C), high-density lipoprotein cholesterol (HDL-C) in the serum were evaluated by FURUNO Clinical Chemistry Analyzer, URUNO ELECTRIC CO., LTD. Nishinomiya, Hyogo, Japan).

### Statistical analysis

Statistical analysis was performed using IBM SPSS software version 25 (IBM Corp., Armonk, NY, USA). Analysis of variance (ANOVA) in conjunction with Tukey’s post hoc test was used to compare among multiple groups, and a difference of p < 0.05 was considered to be statistically significant. The data are presented as the means ± standard error of the mean (SEM).

## Results

### Effects of THC on parameters related to the VHFD-induced obesity mouse model

At the end of the treatment period, autopsy examination revealed that the visceral fat was dramatically increased in VHFD-fed mice compared with LFD-fed mice. The VHFD+THC-treated mice showed that the depots of visceral fat were decreased compared with VHFD-fed mice (Fig 1A).

**Fig 1.**
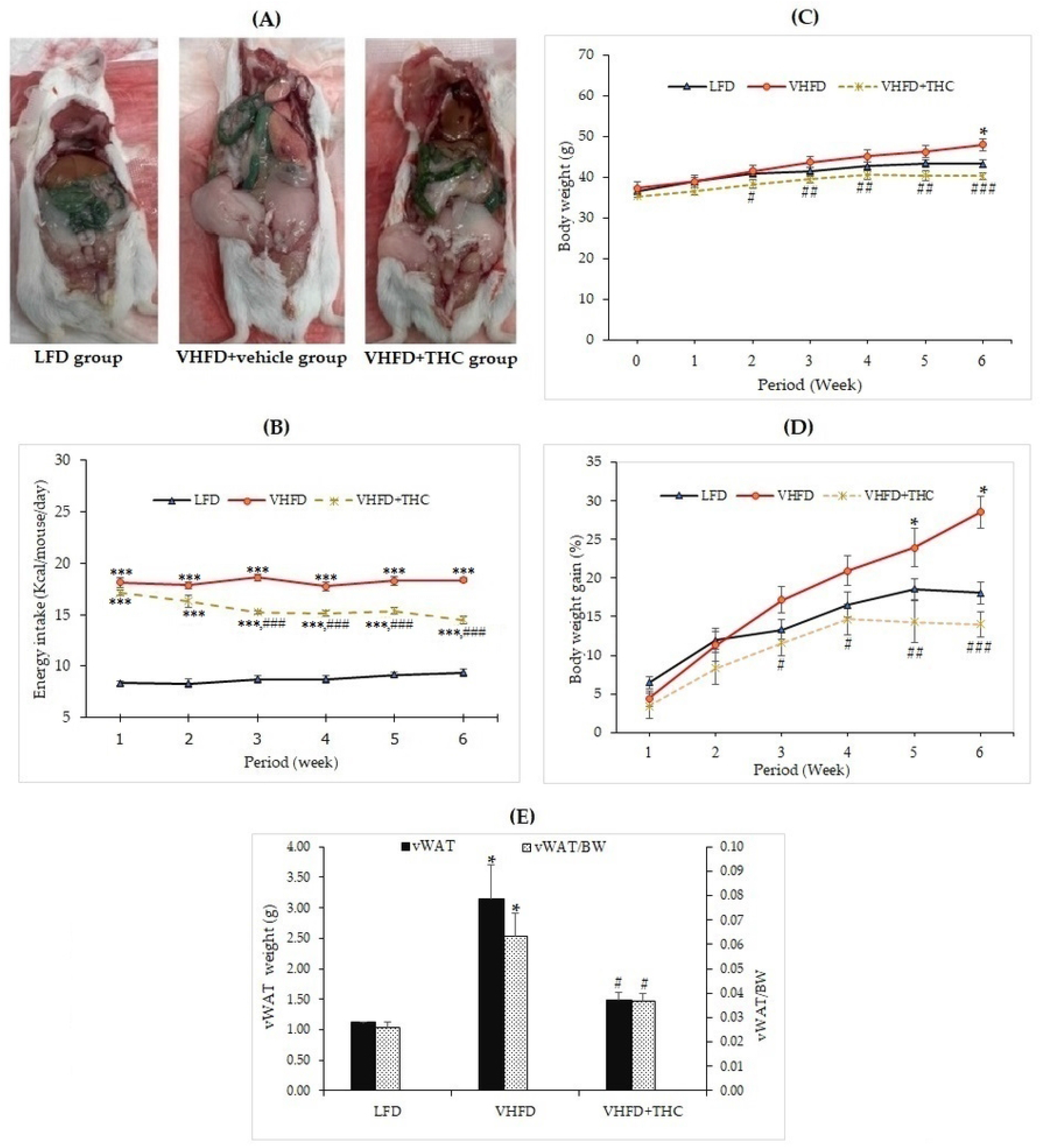
Effects of THC on characteristics of VHFD-induced obese mice: (A) Autopsy examination of adipose tissue distribution after 6 weeks of treatment; (B) Energy intake-time curve; (C) Body weight-time curve; (D) Body weight gain-time curve; and (E) Visceral fat weight and relative adipose tissue. Data are presented as mean ± SEM. ^*^p < 0.05 and ^***^p < 0.001 vs. LFD group; ^#^p < 0.05, ^##^p < 0.005, and ^###^p < 0.001 vs. VHFD group.

The energy intake of the VHFD groups was significantly higher than the LFD group (p < 0.001). Surprisingly, the energy intake was significantly decreased in VHFD-fed mice after 2 weeks of treatment with THC (p < 0.001)(Fig 1B).

The body weight of the mice is presented in Figures 1C and 1D. The untreated VHFD-fed mice displayed significantly higher percent body weight gain and body weight than mice fed with the LFD after 5 and 6 weeks of feeding, respectively (p < 0.05). Interestingly, compared to the VHFD group, VHFD-fed mice treated with THC began showing a significant decrease in body weight starting at 2 weeks post-intervention (Fig 1C) while percent body weight gain significantly decreased after 3 weeks of the treatment with THC and these effects continued until the end of the experimental period (Fig 1D).

The visceral fat weight (VF) significantly increased in the VHFD group compared to the LFD group (p *<* 0.05). THC supplementation for 6 weeks significantly reduced VF weight (p *<* 0.05)(Figure 1E). Moreover, the ratio of the visceral fat to the whole-body mass (relative visceral fat weight/body weight, VF/BW) was significantly increased in VHFD group compared to the LFD group (p < 0.05) (Figure 1E). However, relative adipose tissue was significantly decreased in the VHFD+THC group than those in the VHFD group (p < 0.05), suggesting that THC supplementation prevents the accumulation of the visceral adipose tissue.

### Effect of THC on histological changes of the adipose tissue

Fig 2A shows H&E staining of adipose tissues. The vWATs of the LFD mice exhibited a normal adipocyte structure without any inflammatory cell infiltration. LFD group also displayed normal adipocyte size, whereas the adipose tissue obtained from the VHFD group showed a significant increase in adipocyte size (p < 0.001; Fig 2B) and a large number of CLSs (p < 0.001; Fig 2C). THC supplement significantly lowered the adipocyte size and CLSs compared to the VHFD group (p < 0.001). However, the adipocyte size and CLSs of THC-treated group mice was still higher than that of LFD group (p < 0.001).

**Fig 2.**
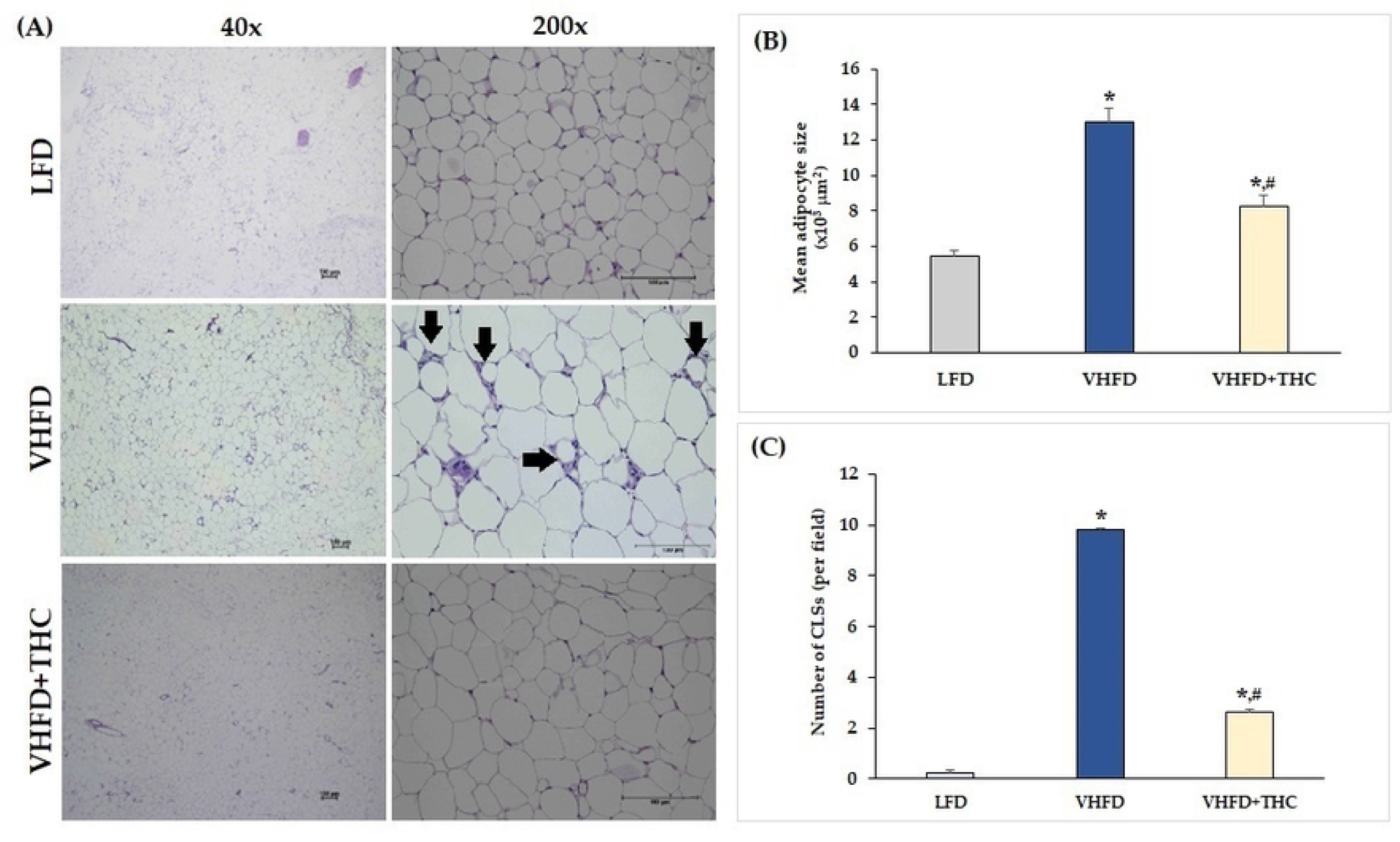
Effects of THC on adipose tissue morphology and inflammatory cell accumulation in VHFD-induced obese mice: (A) H&E staining of vWATs; arrows indicate representative structures of CLSs; left column, magnification 40x; scale bar = 100 μm; right column, magnification 200x; scale bar = 100 μm; (B) Mean adipocyte size; (C) Number of CLSs in WATs. Data are presented as mean ± SEM. ^*^p < 0.001 vs. LFD group; ^#^p < 0.001 vs. VHFD group.

### Effect of THC on serum level of glucose, total cholesterol, HDL, LDL and TG

As shown in Table 2, the levels of serum TG and total cholesterol were significantly higher in VHFD-fed than LFD-fed mice. In addition, the levels of LDL and fasting blood sugar in VHFD group were higher than that in LFD group, albeit no significant difference between groups. Interestingly, THC supplementation significantly reduced blood glucose, total cholesterol, and TG. THC-supplemented mice tended to reduce serum LDL than untreated VHFD mice.

**Table 2.** Blood sugar and lipid profiles.

|  | Group |  |  |
| --- | --- | --- | --- |
|  | LFD-vehicle | VHFD-vehicle | VHFD+THC |
| FBS (mg/dL) | 85.37±6.31 | 102.00±6.49 | 71.63±8.53<sup>#</sup> |
| Cholesterol (mg/dL) | 87.13±4.86 | 104.50±2.23* | 84.75±5.48<sup>#</sup> |
| HDL (mg/dL) | 72.25±3.89 | 69.00±4.74 | 70.50±5.15 |
| LDL (mg/dL) | 8.13±1.01 | 8.25±0.62 | 6.25±0.37 |
| TG (mg/dL) | 107.36±4.15 | 143.25±4.35*** | 96.75±7.22### |

Data are presented as mean ± SEM. \*p < 0.05 and \*\*\*p < 0.001 vs. LFD
group, respectively; #p < 0.05 and ###p < 0.001 vs. VHFD group respectively.

### Effect of THC on adipose angiogenesis

The result showed that the adipose tissue of VHFD-fed mice was highly vascularized, whereas THC-treated mice had lower vascularization (Fig 3A). Furthermore, the microvascular density (MVD), as measured by the percent of CD31 expression, was significantly increased in VHFD-fed group (7.82±0.58) compared to LFD group (4.68±0.48) (p < 0.001). However, a significant reduction of MVD was detected in the adipose tissue of the THC-treated mice (3.29±0.22) as compared with the nontreated VHFD-fed mice (p < 0.001) (Fig 3B).

**Fig 3.**
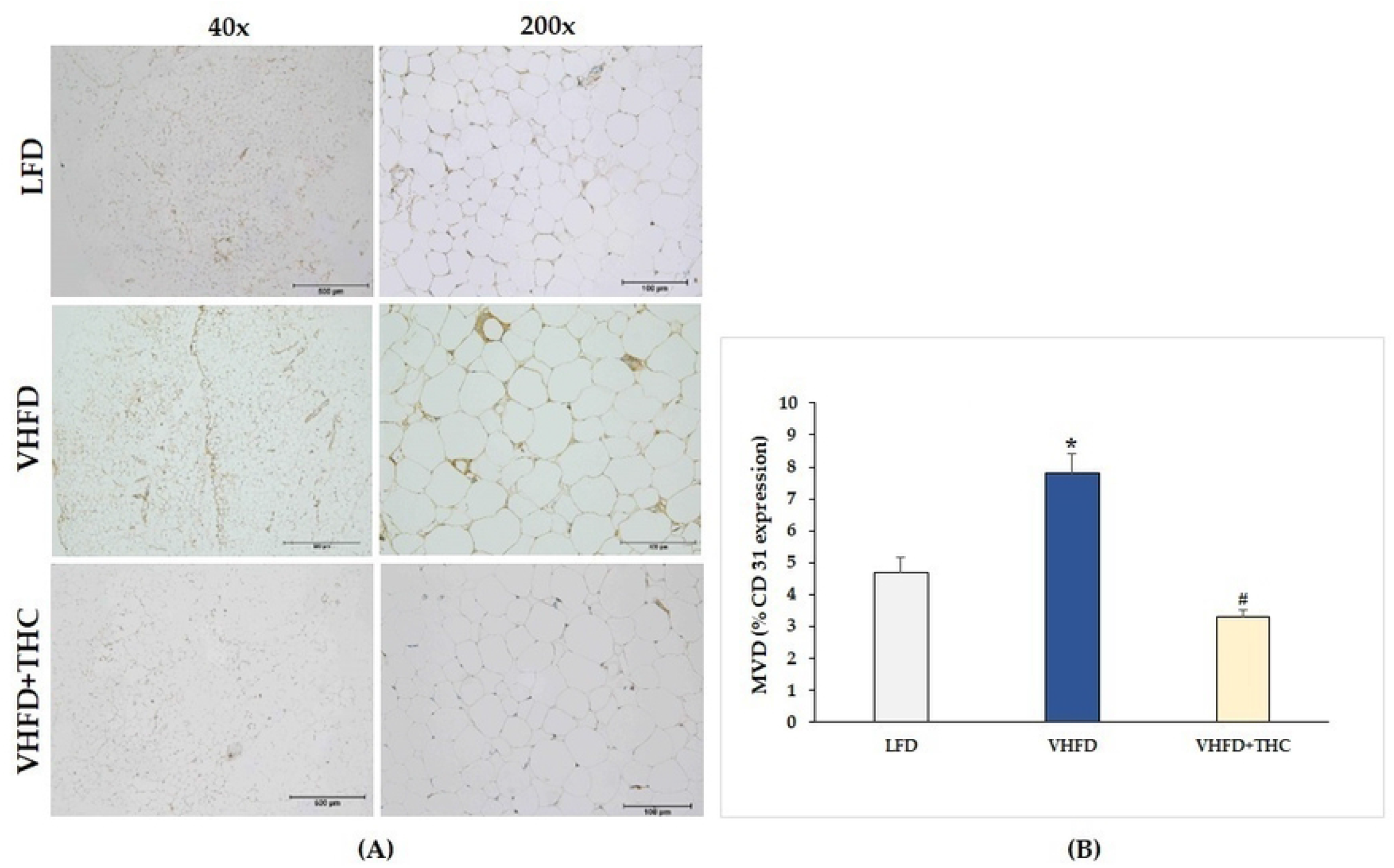
Effect of THC on adipose angiogenesis in VHFD-induced obese mice: (A) Representative CD31 expression; left column, magnification 40×; scale bar = 500 μm; right column, magnification 200×; scale bar = 100 μm; (B) MVD (% CD31 expression). Data are presented as mean ± SEM. ^*^p < 0.001 vs. LFD group; ^#^p < 0.001 vs. VHFD group.

### Effect of THC on adipose angiogenic biomarkers

The use of immunohistochemistry method revealed that the VEGF protein was strongly expressed more in VHFD group than in LFD group. Interestingly, our study demonstrated that THC treated group attenuated VEGF expression (Fig 4A). As shown in Figure 4B, VEGF expression (9.07±0.30%) was significantly higher in the VHFD group than in the LFD group (5.38±0.25%)(p < 0.001). However, significant reductions in % VEGF positive staining were found in THC-treated groups (4.71±0.33)(p < 0.001).

**Fig 4.**
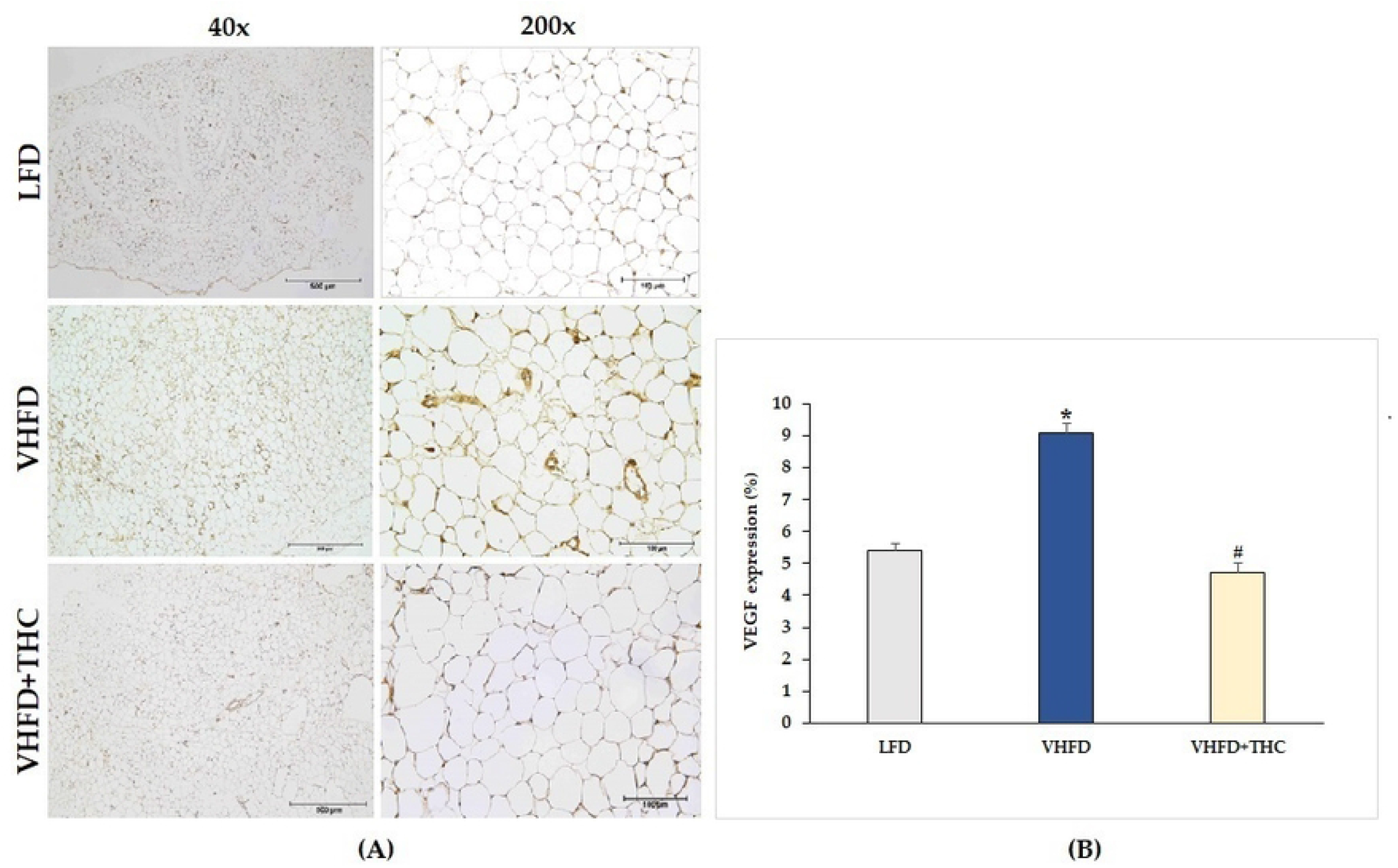
Effect of THC on VEGF expression in VHFD-induced obese mice: (A) Representative VEGF expression; left column, magnification 40×; scale bar = 500 μm; right column, magnification 200×; scale bar = 100 μm; (B) %VEGF expression. Data are presented as mean ± SEM. ^*^p < 0.001 vs. LFD group; ^#^p < 0.001 vs. VHFD group.

The VHFD group also showed stronger VEGFR-2 expression than the LFD group, and the THC treated group showed weaker VEGFR-2 expression than VHFD group (Fig 5A). However, % positive staining intensity for VEGFR-2 expression was not significantly different among groups (Fig 5B).

**Fig 5.**
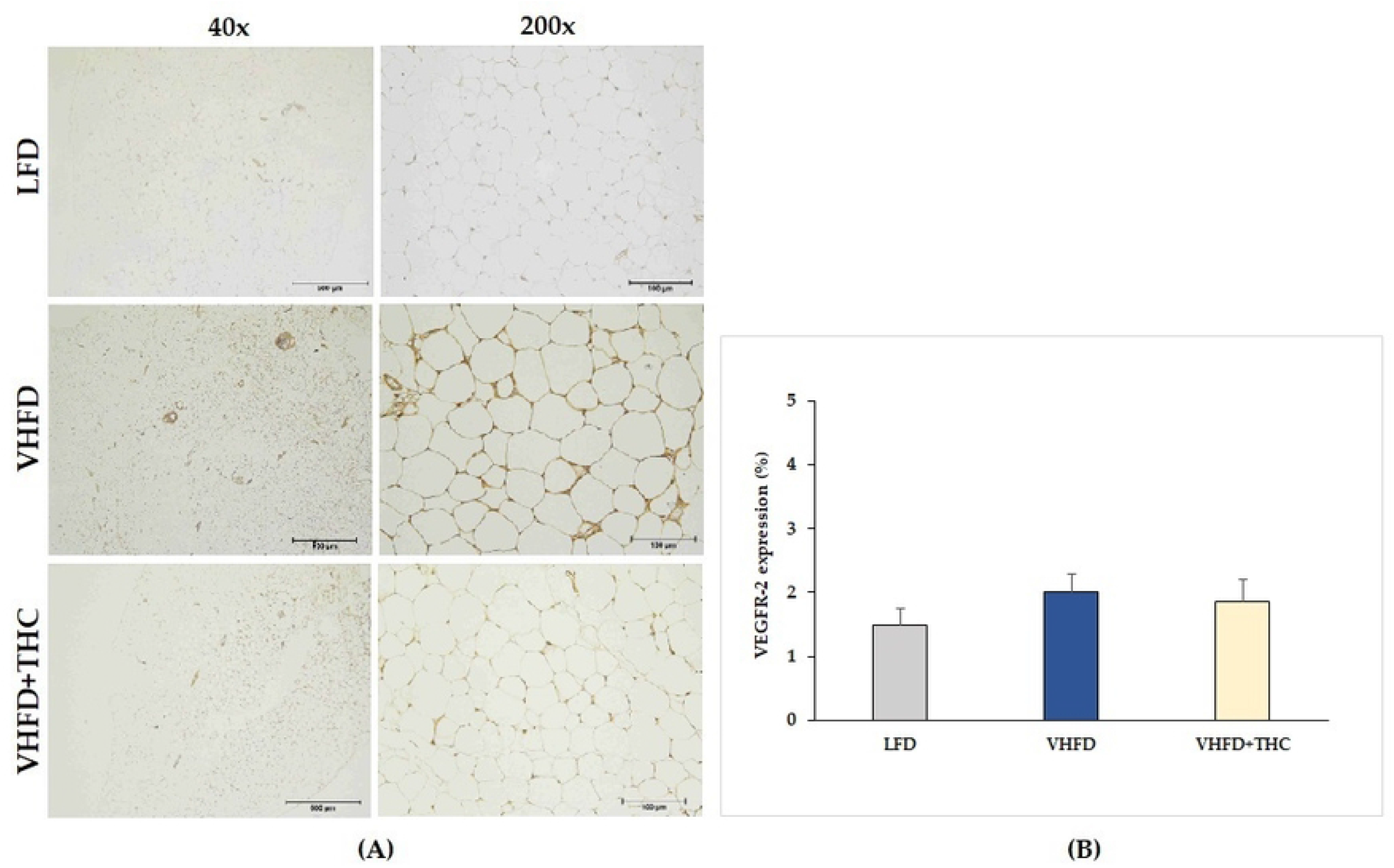
Effect of THC on VEGFR-2 expression in VHFD-induced obese mice: (A) Representative VEGFR-2 expression; left column, magnification 40×; scale bar = 500 μm; right column, magnification 200×; scale bar = 100 μm; (B) %VEGFR-2 expression. Data are presented as mean ± SEM.

### Effect of THC on adipose MMPs expression

MMP-2 and MMP-9 expression dramatically increased in VHFD group, whereas their expression was reduced notably after THC treatment (Fig 6A and 7A, respectively).

**Fig 6.**
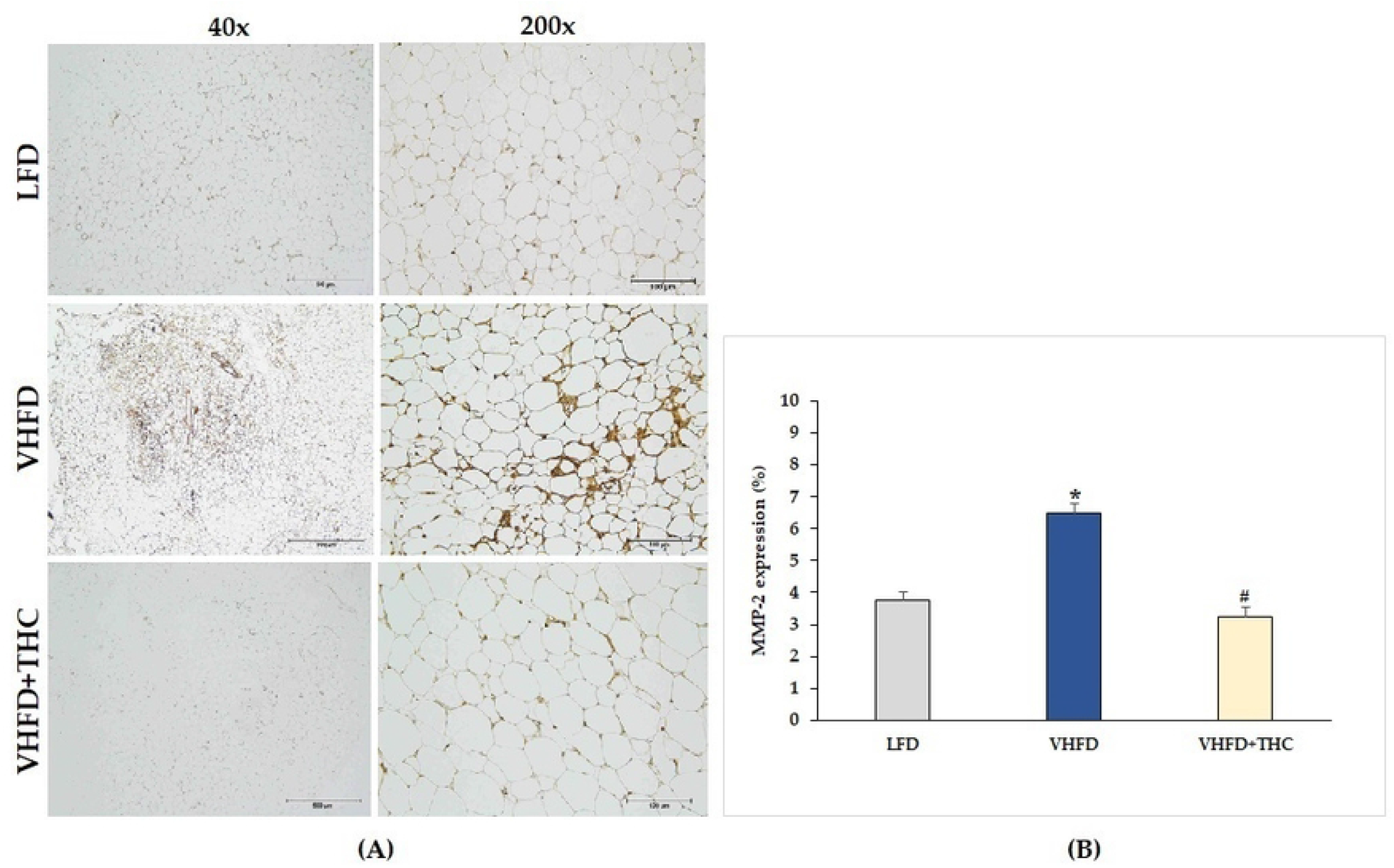
Effect of THC on MMP-2 expression in VHFD-induced obese mice: (A) Representative MMP-2 expression; left column, magnification 40×; scale bar = 500 μm; right column, magnification 200×; scale bar = 100 μm; (B) % MMP-2 expression. Data are presented as mean ± SEM. ^*^p < 0.001 vs. LFD group; ^#^p < 0.001 vs. VHFD group.

**Fig 7.**
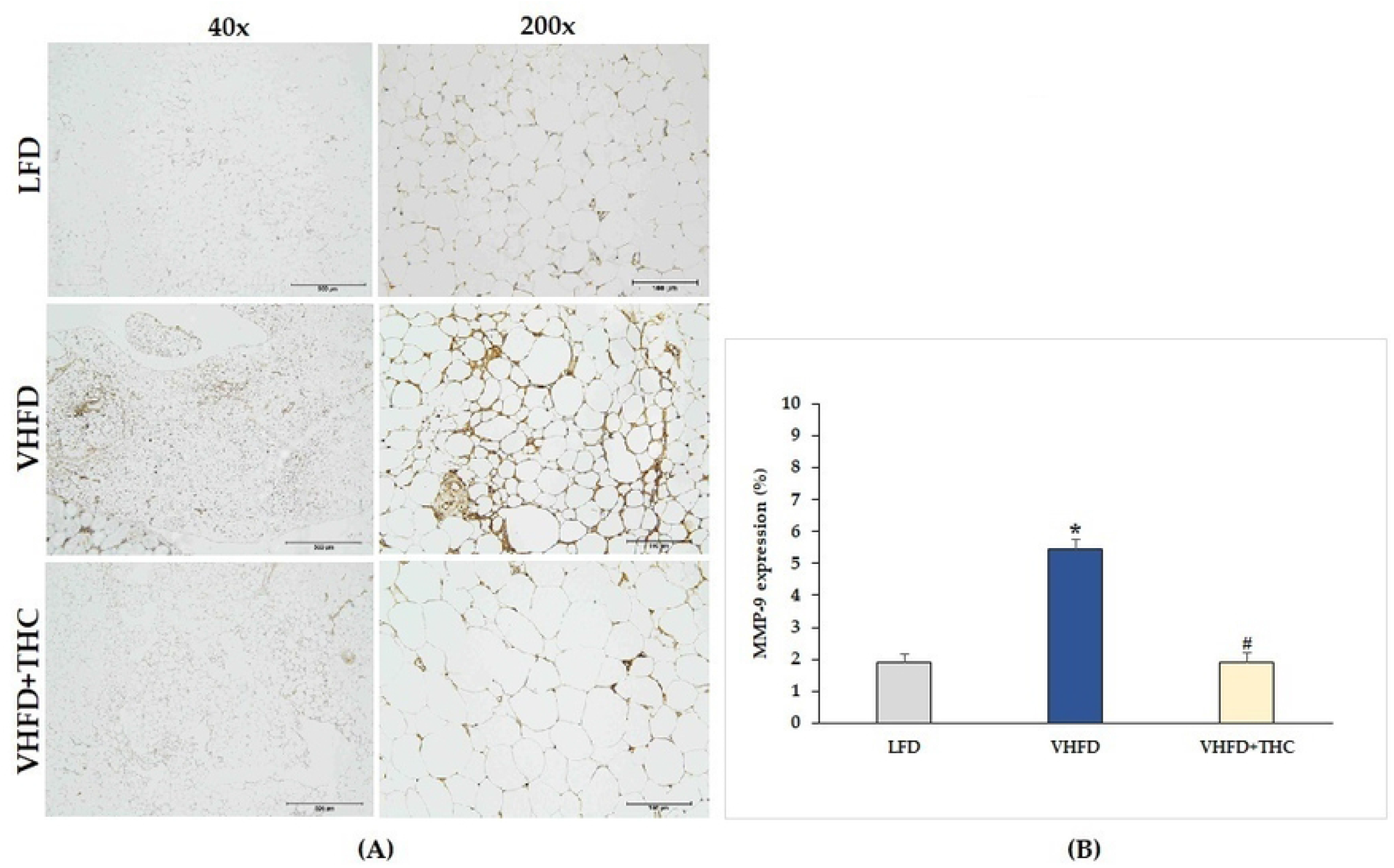
Effect of THC on MMP-9 expression in VHFD-induced obese mice: (A) Representative MMP-9 expression; left column, magnification 40×; scale bar = 500 μm; right column, magnification 200×; scale bar = 100 μm; (B) % MMP-9 expression. Data are presented as mean ± SEM. ^*^p < 0.001 vs. LFD group; ^#^p < 0.001 vs. VHFD group.

Figures 6B and 7B show the percentage of positive staining of MMP-2 and MMP-9, respectively. In the VHFD group, MMP-2 (6.49±0.34%) and MMP-9 (5.44±0.86%) expression were significantly higher than in the LFD group (MMP-2: 3.77±0.18%; MMP-9: 1.92±1.11%)(p < 0.001). Interestingly, in the VHFD+THC group, the percentages of positive staining of MMP-2 (3.24±0.24%) and MMP-9 (1.89±0.64%) were significantly lower than those of the VHFD-vehicle group (p < 0.001).

## Discussion

Due to the faster induction of obesity and stronger metabolic responses, very high fat diet (VHFD) which provides 58–60% kcal fat has recently become a popular alternative to more traditional HFD which provides 40–45% total kcal fat (31). In this study, we used VHFD (60% fat) to induce obesity in the mouse model. We demonstrated significant increases in body weight, body weight gain, visceral fat weight, and relative adipose tissue in the VHFD-fed mice compared with the normal-diet-fed mice (Table 1 and Figure 1). In addition, it was found that the VHFD induced the elevation of triglycerides, blood sugar, total cholesterol, and LDL cholesterol (Table 2). Moreover, VHFD was shown to promote the accumulation of lipids in mature adipocytes, contributing to hypertrophic adipose tissue expansion. In accordance with our previous study, VHFD intake increased body weight, fat accumulation, and adipocyte size in the intra-abdominal compartment, as well as dysfunctional hypertrophic adipocytes. These adipocytes secrete chemokines that recruit immune cells, particularly macrophages, and form CLSs around inflammatory adipocytes, which further promote local and systemic inflammatory responses through the activation of various inflammatory genes including NF-κB p65, MCP-1, TNF-α, and iNOS (30). The VHFD-induced obesity mouse model in the present study demonstrated an increased CLS formation in visceral adipose tissue (Figure 2), indicating the occurrence of the inflammation process, making this model an appropriate *in vivo* model for the evaluation of anti-obesity.

When VHFD-induced obese mice were treated with THC, it was found that THC not only significantly decreased body weight and percent body weight gain but also reduced the visceral fat weight, and relative adipose tissue compared to untreated VHFD-fed mice.

These findings indicate the beneficial effects of THC which can inhibit adipose tissue growth, reduce adipose tissue mass, and prevent body weight gain in obese mice.

Interestingly, there was a significant reduction of food consumption in the VHFD+THC mice compared to the untreated VHFD mice, suggesting that the reduction of body mass gain, visceral fat weight and relative adipose tissue in THC-treated mice might be due to lesser energy intake (Figure 1B).

Oral supplementation with THC in VHFD-fed mice could decrease the levels of serum TG, total cholesterol, fasting blood glucose and tend to decrease LDL cholesterol, suggesting the effect of THC on lipid and glucose metabolism. A previous study also reported that THC significantly reduced body weight as well as delayed adiposity, steatosis, hyperglycemia, and insulin resistance in obese mice (32). They showed that THC markedly alleviated steatosis via the downregulation of lipogenesis, the activation of AMP activated protein kinase (AMPK), and the increase of fatty acid oxidation (32). Moreover, elevated blood glucose and insulin resistance were improved by THC possibly via a regulation of the hepatic insulin signaling cascade, gene transcription involved in glucose metabolism, and reduction of macrophage infiltration in the liver and adipose tissue (32). In the present study, THC was found to significantly reduce inflammatory cell infiltration, as seen by the reduction of CLSs (Figure 2), in vWATs of VHFD-induced obese mice. Overall, this finding emphasizes the effect of THC that can alleviate adipose tissue inflammation, leading to the improvement of metabolic dysfunction including dyslipidemia and hyperglycemia.

Obesity progression has demonstrated links with angiogenesis and remodeling of extracellular matrix (ECM) (33). By interfering with the angiogenesis process, the hypertrophy and hyperplasia of adipose tissue should be subsided. In the present study, the microvascular density in adipose tissue was examined by using an anti-CD31 antibody. As expected, adipose tissue of mice fed with VHFD was found to be highly vascularized, and the greater vascular density of visceral adipose tissue, the higher expression of VEGF was found.

VEGF is an important factor in angiogenesis, and visceral adipose tissue has been shown to have the expression of VEGF at the highest level (34, 35). Endothelial cells along with infiltrated inflammatory cells and stromal cells of adipose tissues contribute to VEGF production. The VEGF families currently includes VEGF-A, -B, -C, -D, -E, -F, and PlGF (placental growth factor), which bind in a distinct pattern to three structurally related receptors namely VEGFR-1, -2, and -3 (13). The VEGF-A/VEGFR-2 pathway plays a central role in angiogenesis. Ejaz *et al*. demonstrated that supplementing the high-fat-diet of mice with curcumin could lower body weight gain, the growth of adipose tissue, and angiogenesis in the adipose tissue thru the reduced expression of VEGF and VEGFR-2 (15).

In the present study, it was found that supplementing the VHFD with THC 300 mg/kg for 6 weeks markedly reduced microvascular density in adipose tissue along with reduced expression of VEGF. The results are remarkable considering the rather short duration (6 weeks) compared to the previous report by Ejaz *et al* (15). At the dose of 300 mg/kg daily, THC has been shown to possess an anti-angiogenetic activity in hepatocellular carcinoma in mice (29). It is noteworthy that in the present study, the percentages of VEGFR-2 expression were also higher in VHFD-fed mice than the controls, and THC treatment tend to decrease VEGFR-2 expression, but these differences were not statistically significant. These findings suggest that the anti-angiogenic effect of short-term THC supplement was due to the downregulation of VEGF expression but not VEGFR-2 expression.

Besides vascular growth factors, adipocytes release several MMPs which play a key role in angiogenesis. Adipose-tissue MMPs including the two major MMPs (MMP-2 and MMP-9) could potentially affect preadipocyte differentiation and microvascular maturation by modulating ECM (33). Moreover, MMP-9 can release the matrix bound VEGF, resulting in the neo-vascularization process (36). In this study, increased expression of MMP-2 and MMP-9 was found in the adipose tissues of VHFD-fed mice. Interestingly, THC-treated mice had a significant decrease of MMP-2 and MMP-9 expression, suggesting that THC inhibits adipose angiogenesis via the reduction of MMP-2, MMP-9 as well as VEGF expression.

## Conclusions

The present study highlights the short-term beneficial effects of THC (the major metabolite of curcumin) that can suppress angiogenesis in adipose tissue via the reduction of VEGF, MMP-2, and MMP-9 expression in the VHFD-induced obese mice model. This anti-angiogenic activity, together with the suppression of food consumption, appear to be responsible for the lower body weight gain, visceral fat weight, relative adipose tissue, and, ultimately, the alleviation of dyslipidemia and hyperglycemia in the obese mice. Overall, our study demonstrated the potential benefit of THC in mitigating obesity and associated metabolic disorders as well as elucidating the suppression of angiogenesis in adipose tissue as one of its underlying mechanisms.

## Funding

This research was funded by the Research Fund of Faculty of Medicine (2-19/2665), the Thailand Science Research and Innovation Fund fiscal year 2023 (TUFF51/2566), and the Research Group in Exercise and Aging-Associated Diseases, Faculty of Medicine, Thammasat University, Thailand.

## Data Availability Statement

The animal study protocol was approved by the Animal Care and Use Committee of Thammasat University, Thailand (ethical approval code 020/2021).

## Conflicts of Interest

The authors declare no conflict of interest.

## Notes

### Competing Interest Statement

The authors have declared no competing interest.

